# On-Demand Expansion Fluorescence and Photoacoustic Microscopy (ExFLPAM)

**DOI:** 10.1101/2023.07.19.549787

**Authors:** Xuan Mu, Chenshuo Ma, Xuan Mei, Junlong Liao, Rebecca Bojar, Sizhe Kuang, Qiangzhou Rong, Junjie Yao, Yu Shrike Zhang

## Abstract

Expansion microscopy (ExM) is a promising technology that enables nanoscale imaging on conventional optical microscopes by physically magnifying the specimens. Here, we report the development of a strategy that enables i) on-demand labeling of subcellular organelles in live cells for ExM through transfection of fluorescent proteins that are well-retained during the expansion procedure; and ii) non-fluorescent chromogenic color-development towards efficient bright-field and photoacoustic imaging in both planar and volumetric formats, which is applicable to both cultured cells and biological tissues. Compared to the conventional ExM methods, our strategy provides an expanded toolkit, which we term as expansion fluorescence and photoacoustic microscopy (ExFLPAM), by allowing on-demand fluorescent protein labeling of cultured cells, as well as non-fluorescent absorption contrast-imaging of biological samples.

## Introduction

Expansion microscopy (ExM) has emerged as a promising technology for observing biological samples since it enables nanoscale imaging on conventional optical microscopes.^1-4^ The key of ExM is the physical magnification of the specimens, where a swellable hydrogel plays an important role during this process.^5,6,7^ In general, for ExM, target biomolecules and fluorescent tags are covalently anchored to the hydrogel network to maintain the relative position in the hydrogel.^8,9^ Specific proteolytic enzymes are introduced to break the cellular skeleton to avoid heterogeneous hydrogel swelling.^10,11^ Finally, the hydrogel along with the specimen is expanded by immersing in a liquid of low osmolarity, which leads to an amplification of ∼4.5× in linear dimension in a typical setup.^2^ Up to now, various ExM methodologies have been developed for nanoscale imaging of proteins and nucleic acids within cells,^12,13^ which has already enabled successful high-resolution characterizations of samples that are both small, planar as well as large, three-dimensional, spanning across bacterial cells,^14^ mammalian cells,^15^ tissue sections,^16^ tissue blocks,^17^ and whole organisms.^18^

Many efforts have been made to enhance the resolution limitation of ExMs, such as using hydrogels with an increased swelling ability or repeating the expansion process,^19,20,21^ so far improved the magnification to ∼20×.^22^ To further enhance the ExM imaging, additional focus has been placed on the development of labeling strategies that ensure high labeling densities.^23,24,25^ For such a perspective, genetically encoded fluorescent proteins through transfection might be a good alternative to antibodies. Similar to immunostaining, transfection methods can generate bright and photostable fluorescent signals, leading to convenient visualization of the labeled structures without the need for additional staining steps.^26,27^ In addition, transfection methods may now be processed with ready-to-use reagents, eliminating complex protocols.^28-31^ Nevertheless, on-demand transfection applied towards ExM usage has not been demonstrated yet.

Given the fact that fluorescent proteins have been shown to be directly expandable while maintaining their signals post-expansion,^12,32^ we therefore reasoned that transfection-enabled rapid labeling could potentially be applied to ExM as well. To confirm the hypothesis, we selected fluorescent proteins (green fluorescent protein, GFP and red fluorescent protein, RFP) to label subcellular organelles due to their resistance to proteinase-based digestion during the expansion process (**Figure 1**).^33,34^ Since Proteinase K is required for digestion to make samples mechanically homogenized, it is important to use proteinase-resistant proteins to preserve fluorescence signal.

**Figure 1.**
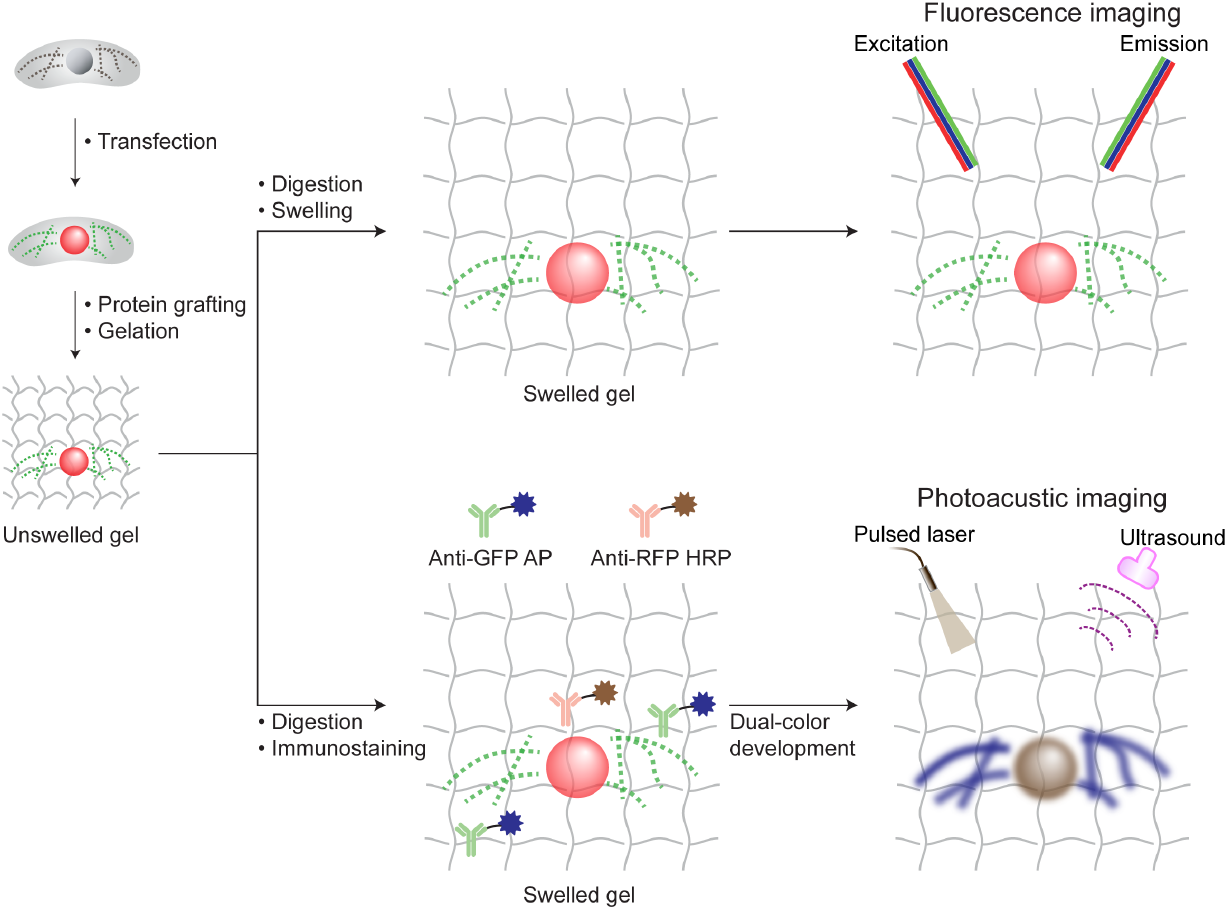
Design principle of transfection-enabled ExFLPAM. Cellular organelles, such as mitochondria and nuclei, can be labeled by the transfection of fluorescent proteins (e.g., GFP and RFP), followed by grafting GFP and RFP onto the gel network. GFP and RFP are resistant to proteinase-based digestion. After digestion and swelling, the samples can be imaged under fluorescence microscope with improved spatial resolution. In addition, GFP and RFP can be further immunolabelled with enzymes such as AP and HRP. The swollen gel, after adding enzyme substrates, would develop visual colors, which may then be imaged by photoacoustic microscopy with volumetric imaging capability and relatively large imaging depth.

With fluorescence microscope, the expanded samples were confirmed to show details in these labeled subcellular structures. In the meanwhile, to further broaden the suitability of the labeled specimens for a wide range of optical imaging modalities beyond fluorescence imaging, GFP and RFP were additionally immunolabelled with alkaline phosphatase (AP) and horseradish peroxidase (HRP), respectively. When subsequently incubated with Vector Blue and NovaRED, the enzyme substrates to AP and HRP, the structures produced corresponding blue and reddish-brown chromogen precipitants following the reactions. The chromogen precipitants, although nonfluorescent, have developed visual colors with strong optical absorption, which can then be imaged by photoacoustic microscopy with favorable volumetric imaging ability and advanced imaging depth. Our methodology provides a new scheme to rapidly label subcellular organelles of interest for ExM, which also for the first time enables both fluorescence imaging and photoacoustic imaging with extended imaging capacities.

The feasibility of the standard ExM protocol^2^ was first confirmed on multiple cell types before the introduction of our intended transfection method. As shown in **Figure S1a**, C2C12 myoblasts were immunostained with fluorescent antibody against tubulin and then processed with expansion. As anticipated, a ∼4.3-fold linear expansion in tubulin diameter was achieved (**Figure S1b**), which led to more sharply resolved microtubules than could be observed before expansion.

To subsequently confirm the compatibility of ExM with on-demand transfection-enabled labeling of subcellular organelles, wildtype cells without any intrinsic fluorescence emission were transfected with CellLight regents for 24 h immediately prior to expansion, to explicitly express fluorescent proteins of desired colors on target structures. In one demonstration, the nuclei of MCF-7 cells were labeled with RFP. After treating the samples with the ExM protocol followed by swelling in 1× PBS and then water, the nuclei showed significantly improved detail in structures compared to non-expanded samples (**Figure 2a**). As shown in **Figure 2b**, in another dual-color demonstration, the mitochondria of MCF-7 cells were labeled with GFP while the nuclei were transfected to express RFP. With expansion, both mitochondria and nuclei were clearly expanded with substantially improved resolutions under fluorescence imaging. The expansion ratios of both organelles were quantified, resulting in ∼1.7-fold in 1× PBS and ∼3.1-fold in water, with no noticeable differences between the two labeled organelles (**Figure 2c**) or with single labeling of the nuclei (**Figure 2a**). Different from conventional immunostaining, transfection-induced fluorescent protein expression would be present and fused to the localization sequence within targeted intracellular structure before expansion. The fusion of proteinase-resistant fluorescent proteins (GFP and RFP) may protect these original proteins from digestion and result in insufficient mechanical homogenization. Thus, the expansion ratio reduced slightly comparing to the original expansion protocol.

**Figure 2.**
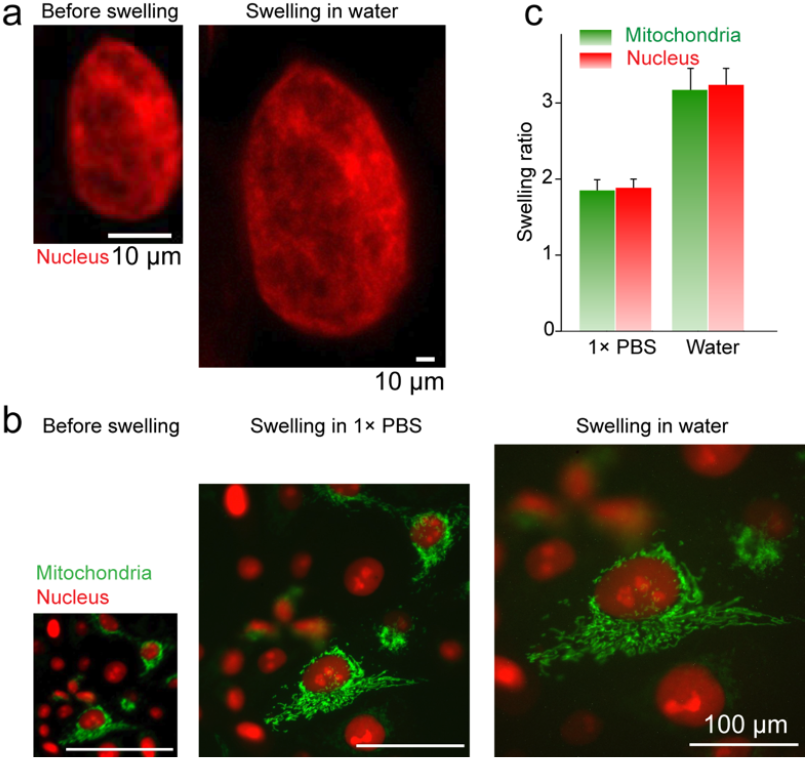
On-demand transfection-enabled ExM. a. Fluorescence imaging of the same RFP-labeled MCF-7 nucleus before and after swelling in water. b. Fluorescence imaging of MCF-7 cells on-demand transfected for 24 h prior to ExM, showing GFP-labeled mitochondria and RFP-labeled nuclei in three different swelling statues after transfection (without swelling and swollen in 1× PBS and water). c. Quantitative characterization of the swelling ratios of both organelles in 1× PBS and water, compared to that without swelling.

While fluorescence imaging replies on the fluorescent emission of the transfected proteins, photoacoustic microscopy (PAM) does not. In fact, by detecting the optical absorption contrast, PAM can image both fluorescent and non-fluorescent molecules, which is much more flexible than traditional fluorescence imaging. In PAM, pulse-laser light is absorbed by biomolecules and the absorbed photon energy is converted into heat via the photothermal effect. The heat-induced expansion generates acoustic pressure waves that can be detected to map the original optical energy deposition in the sample. PAM is intrinsically a volumetric imaging modality, and takes advantage of the rich optical contrast of endogenous or exogenous biomolecules, as well as the deep-penetrating acoustic detection for superior imaging depth. We have previously demonstrated that PAM is able to perform multi-spectrum imaging of non-expanded biological samples stained with color-absorbing histology dyes.^35^ To this end, we rationed that, not only is it possible to retain the fluorescence signals post-expansion, but these fluorescent proteins may be further converted to optical absorption contrast for PAM, by applying, further, a color-developing protocol that is commonly used in conventional histology, which we term as ExFLPAM.

To be imaged by the PAM, the specimens must carry optical absorption contrasts for pulsed laser excitation and ultrasonic detection.^36,37^ In an initial demonstration, a dual-absorbing-color staining strategy was developed by using chromogenic substrates for immunolabeled enzymes on fluorescent proteins. As examples, GFP-HUVECs were cultured, fixed, and immunolabeled with AP-anti-GFP; when they were expanded in 1× PBS and subsequently exposed to Vector Blue substrate specific for AP, strong blue-colored precipitations were produced leading to the entire gel turning blue visually (**Figure 3a**). Similarly, when RFP-HUVECs were labeled with HRP-anti-RFP, expanded, and treated with NovaRED substrate specific to reaction with HRP, a strong brown-colored chromogenic contrast was developed (**Figure 3b**). Since these are absorbing-only contrasts, the cells were visible under bright-field microscopy without the need for fluorescence imaging, indicating blue and reddish-brown colors, respectively (**Figure 3c, d**).

**Figure 3.**
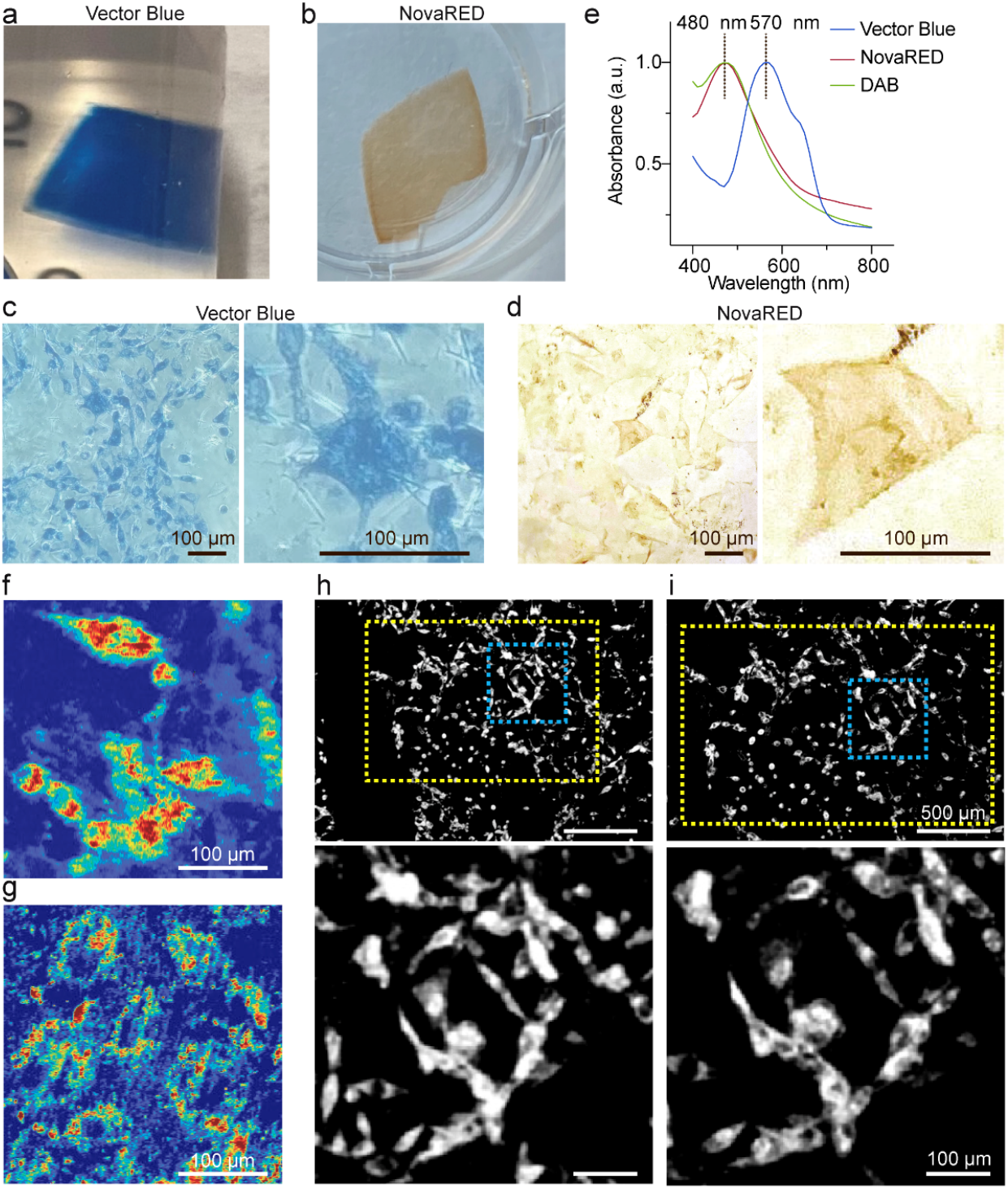
Color staining and photoacoustic imaging of cell samples. a. Vector Blue, a substrate of AP, developed blue color for AP-labeled cell samples. b. NovaRED, a substrate of HRP, developed brown color for HRP-labeled cell samples. Transmission-mode optical microscope image of (c) blue- and (d) brown-stained HUVEC (transfected with GFP and RFP, respectively) in the gel. E. Absorbance spectra of the stained color by Vector Blue, NovaRED and DAB. Photoacoustic imaging of stained HUVECs in the gel at 532 nm, f. Vector Blue; g. NovaRED. PAM images of Vector Blue gel before (h) and after (i) swelling in water. Yellow dotted squares indicate the same area before and after swelling. Bottom row: close-up images of selected areas within blue dotted boxes.

As previously discussed, the precipitations formed by these chromogens used for staining have strong optical absorptions across a wide wavelength range. **Figure 3e** shows typical absorption spectra of precipitations formed from Vector Blue, NovaRED, and ImmPACT DAB substrates upon reaction with their corresponding specific enzymes. Accordingly, we developed a high-resolution PAM system with a spatial resolution of ∼600 nm (**Figure S2**). It was clearly shown that the expanded cells stained with both Vector Blue (**Figure 3f**) and NovaRED (**Figure 3g**) could be imaged at 532 nm. The expansion ratio was also calculated on the Vector Blue-stained samples. Compared to cells before expanding (**Figure 3h**), the gel expanded 1.32-fold to 1.37-fold after immersion in water (**Figure 3i**). The expanded gel was further incubated in 10× PBS, where a shrinkage of 1.68-fold to 1.78-fold was observed (**Figure S3**). The imaging of expanded and shrunken specimens was further confirmed with additional PAM images (**Figure S4**). Due to the use of 1× PBS during the labeling process, these samples were already slightly expanded prior to immersion in water. As compared to the transfected-only samples (**Figure 2**), the expansion ratio was reduced from 1.82-fold to 1.37-fold after chromogenic labeling, which was possibly due to the formation of the precipitates on the hydrogel networks during the enzymatic reactions of chromogens that mechanically limited the full expansion of these gels. A similar observation was made when the expanded gel was shrunken back in 10× PBS. The NovaRED-stained cells exhibited a similar expansion behavior for PAM imaging (**Figure S5**). Notably, taking advantages of the acoustic penetration of PAM, volumetric cellular imaging was acquired from 0 μm (surface) to 30 μm in depth both before and after expansion (**Figure S6**).

Finally, based on the above results, the ExFLPAM of mouse kidney tissues expressing GFP were further tested using the same absorbing color-enabled staining strategy. As revealed in **Figure 4a**, the whole kidney slice could be successfully imaged by PAM at the wavelength of 532 nm, post-expansion in 1× PBS and post-staining with Vector Blue. Again, with the volumetric imaging ability of PAM, the expanded, stained tissue at different depths (0, 25 and 50 μm) was readily imaged (**Figure 4b**). The selected areas in **Figure 4b** were further imaged at a higher resolution, showing detailed kidney structure such as convoluted proximal tubules (as indicated with arrows in **Figure 4c**) otherwise not resolvable by PAM without expansion.

**Figure 4.**
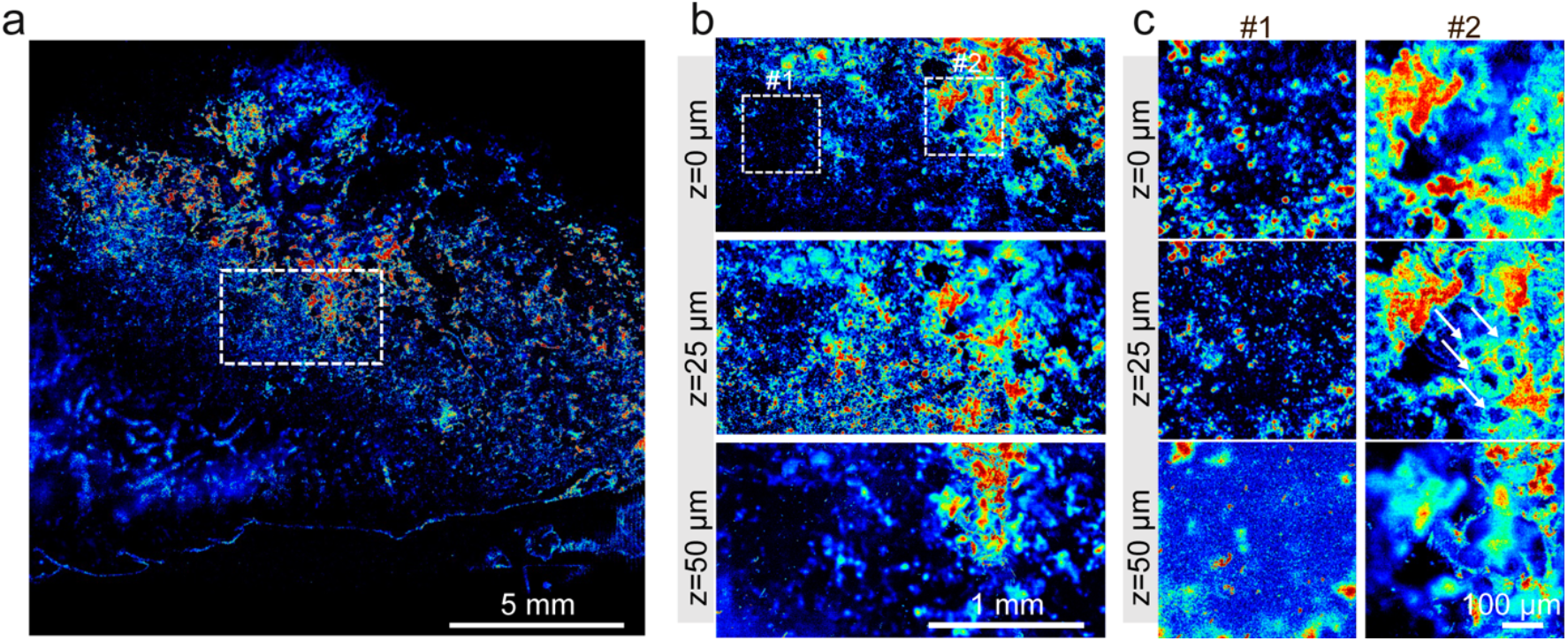
ExFLPAM for kidney slices. A. Whole view of the kidney sample. B. Select areas with different depths. C. Selected areas #1 and #2 with different depths.

In summary, we have developed a new strategy that enables i) on-demand labeling of subcellular organelles in live cells for ExM through transfection of fluorescent proteins that are well-retained during the expansion procedure; and ii) chromogenic color-development towards efficient bright-field and PAM imaging in both planar and volumetric formats, which is applicable to both cultured cells and biological tissues. These advances will likely further expand the applicability of ExM leading to additional capacities not possible or convenient before using immunostaining-based fluorescence imaging or fluorescent protein-labeled structures that does not allow absorbing contrasts.

Nevertheless, our proof-of-concept studies do not come with no limitations and warrant further developments. For example, to simplify the conceptual demonstrations, we sticked to the very original yet most robust expansion protocol that only provides an expansion ratio of ∼4-fold,^2^ which in certain cases such as transfection, was further reduced to ∼3-fold possibly due to the insufficient mechanical homogenization after the enhanced expression of proteinase K resistant protein (GFP and RFP). The expansion ratio was also reduced after chromogenic labeling, which may be due to the confinement of the precipitates formed with the chromogen under enzymatic reactions, on the hydrogel network. Optimizations that would further improve the expansion ratio shall be investigated. Moreover, compatibility of such a concept with other ExM variations, for example, iterative ExM,^17,20^ remains to be explored. Multi-spectral PAM imaging of multiple contrasts at the same time could be explored as well,^38^ given the broad availability of these chromogens that all have distinct absorption spectra across the wavelength range (e.g., **Figure 3e**). We anticipate that, with additional technical advancements based on our current concept, the strategy will find potential wide applicability in biology, bioimaging, biomedicine, and beyond.

## Acknowledgements

We acknowledge the support by the National Institutes of Health (R03EB122662) and the Brigham research Institute.

