## Supplement Information for "On-Demand Expansion Fluorescence and Photoacoustic Microscopy (ExFLPAM)"

### Methods

#### Cell culture

For MCF-7 cells, the following cell medium was used: Dulbecco's modified Eagle medium (DMEM; 21063029, Gibco), supplemented with 10% fetal bovine serum (FBS; 10438026, Gibco), and 1% mL $L^{-1}$ penicillin-streptomycin (15140122, Gibco). For GFP-prelabeled or RFP-prelabeled HUVECs, Endothelial Cell Medium (1001, ScienceCell) was used. The cells were cultured at 37 °C with 5% $CO_2$. The cells were detached from flasks using 0.05% trypsin-ethylenediaminetetraacetic acid (EDTA) (25300054, Gibco), centrifugated and diluted to 50,000 cells $mL^{-1}$. For 24-well plate, a coverslip (13 mm in diameter) was inserted into each well and 1 mL of diluted cell suspension was added. The cells were cultured for 24 h before further use.

#### Cell fixation

For cell fixation, cytoskeleton extraction buffer was prepared, which included Triton X-100 (X100, Sigma, 5% (w/v), 4 mL), 4-piperazinediethanesulfonic acid (PIPES; P6575, Sigma-Aldrich 1 M, pH adjusted to 7.0 with 5-M sodium hydroxide solution, 4 mL), ethylene glycol-bis(2-aminoethylether)-N,N,N,N-tetraacetic acid (EGTA; 03777, Sigma-Aldrich, 10 mM, 4 mL), magnesium chloride (AM9530G, Invitrogen, 40 µL), and water (27.96 mL). For microtubule fixation, a fixation solution was prepared as following: 16% (w/v) paraformaldehyde (158127, Sigma-Aldrich, 1.5 mL), 8% (w/v) glutaraldehyde (G6257, Sigma-Aldrich, 0.1 mL), PBS 10× (0.8 mL), and water (5.6 mL). A reduction solution was prepared by dissolving sodium borohydride (452882, Sigma-Aldrich, 10 mg) in PBS (10 mL). A quenching solution was prepared by using glycine (G7126, Sigma-Aldrich, 375 mg), PBS 10× (5 mL), and water (45 mL).

For each well, cell culture medium was replaced by 500 µL of cytoskeleton extraction buffer and incubated 1 min at room temperature. Then the cytoskeleton extraction buffer was aspirated and 1 mL of microtubule fixation solution was added. After incubating for 10 min at room temperature, cells were treated with reduction solution for 7 min and with quenching solution for 10 min, separately.

#### Immunostaining

To perform immunostaining, the samples were treated with 300 µL of MAXblock (15252, Active Motif) blocking medium for 2 h, followed by washing with 1 mL of MAXwash washing medium

(15254, Active Motif) for 10 min. The cells were incubated with the primary antibodies for 2 h at room temperature and another 2 h with secondary antibodies in the dark. The following primary antibodies were used: sheep polyclonal anti-tubulin (ATN02, Cytoskeleton, 1:500) and rabbit polyclonal anti-tubulin (AB6046, Abcam, 1:100). The secondary antibodies used were listed as below: goat anti-rabbit Alex 564 (A11035, Invitrogen, 1:100), donkey anti-sheep Alex 546 (A21098, Invitrogen, 1:100), rabbit anti-sheep HRP-conjugated (AB6747, Abcam, 1:500), anti-GFP (GOAT) antibody ALP-conjugated (600-105-215, Rockland Immunochemicals, 1:200) and anti-RFP (RABBIT) antibody HRP-conjugated (600-403-379, Rockland Immunochemicals, 1:200).

**Transfection**

For cell transfection, the following CellLight™ regents were used. CellLight™ Mitochondria-GFP (C10600, Invitrogen) and CellLight™ Nucleus-RFP (C10603, Invitrogen). According to manufacturer's instruction, cells were incubated with transfection regents for 24 h before imaging.

**Gelation and digestion**

To prepare the AcX stock solution, 5 mg of acryloyl-X (A20770, Invitrogen) and 500 μL of anhydrous DMSO (D12345, Invitrogen) were mixed. The stock solution was diluted in PBS (1:100) before use. To prepare the gelation solution, stock X (98%), TEMED (T7024, Sigma, 1%), and APS (248614, Sigma, 1%) were mixed on ice before use. The stock X was composed as following: sodium acrylate (408220, Sigma-Aldrich, 3.8 g per 10 mL), 0.5 mL of acrylamide (A9099, Sigma-Aldrich, 0.5 g $mL^{-1}$), 0.75 mL of N,N-methylenebisacrylamide (146072, Sigma-Aldrich, 0.02 g $mL^{-1}$), 4 mL sodium chloride (5 M, Sigma-Aldrich, 14.61 g per 50 mL,), 1 mL PBS 10×, and 1.3 mL water.

The AcX solution was diluted in PBS (1:100 dilution) and added to cells for incubating overnight. After washing with PBS for 5 min on ice, 300 μL of gelation solution was added to each well and incubated for 5 min on ice. A customized gelation chamber was made according to previous study.[1] The cell culture coverslips were inverted and contacted with 40 μL of gelation solution in the gelation chamber. After incubated at 37 °C for 1 h, the gelation solution was. The cell culture coverslips with gel were collected from the chamber and transferred to glass slides. Strong digestion was performed by using strong digestion solution mixed with 99% digestion buffer and 1% proteinase K (P8107S, NE Biolabs). Strong digestion solution was prepared as follow: 10 mL of Triton X-100 (T8787, Sigma-Aldrich, 5 w/v%), 0.2 mL of EDTA (15575020, Invitrogen), 5 mL of tris(hydroxymethyl)aminomethane (Tris; AM9856, Invitrogen), 20 mL of sodium chloride (S6546, Sigma-Aldrich). For digestion, samples were kept on a shaker with digestion solution at room temperature overnight in the dark.

**Color development through chromogenic staining**

As described before, GFP-HUVECs were cultured, fixed, immunolabeled with AP-anti-GFP (600-105-215, Rockland Immunochemicals, 1:200) and expanded in 1× PBS. For color development, samples were then exposed to Vector Blue substrate specific for AP (SK-5300, Vector Laboratories). To prepare the substrate working solution, Vector Blue Reagent 1 (80 μL), Vector Blue Reagent 2 (80 μL) and Vector Blue Reagent 3 (45 μL) were added to Tris-HCl buffer (AM9855G, Invitrogen, 5 mL, pH=8.2-8.5). The samples were incubated with the substrate working solution for 30 min to develop the color and washed with 1× PBS for 5 min before imaging.

RFP-HUVECs were treated following the same protocol but immunolabeled with HRP-anti-RFP (600-403-379, Rockland Immunochemicals, 1:200). NovaRED kit (SK-4805, Vector Laboratories) was further used for color development. A substrate working solution was prepared according to the manufacturer's instruction. Briefly, NovaRED Reagent 1 (80 μL), NovaRED Reagent 2 (50 μL), NovaRED Reagent 3 (50 μL) and NovaRED Reagent 4 (80 μL) were added to NovaRed Diluent (5 mL). The samples were incubated with the substrate working solution for 15 min at room temperature and washed with 1× PBS for 5 min.

**Expansion**

After digestion, the detached gel was carefully collected. 10 mL of water was added to the sample and incubated at room temperature for 10 min for expansion. The expansion process was repeated 3 times. In other cases, PBS of different concentrations were also used for expansion.

**TOR-PAM**

TOR-PAM employs a tightly focused laser beam that is diffraction-limited, with the lateral resolution being primarily determined by the optical focal spot size. A laser (VPFL-G-20, Spectral Physics) was operated at 532 nm to provide the excitation light. The laser beam was focused by an objective lens (Mitutoyo 50x M Plan APO) with an NA of 0.55, and delivered to the sample surface with a pulse energy of 800 nJ at 532 nm. On the other side of the sample, an ultrasonic transducer with a central frequency of 30 MHz (V214-BB-RM, Olympus-NDT) was used to detect the resultant photoacoustic signals. The sample was excited with LED light from the top, which after passing through the sample, was reflected in another direction by a dichroic mirror. An image was captured by a CCD camera.

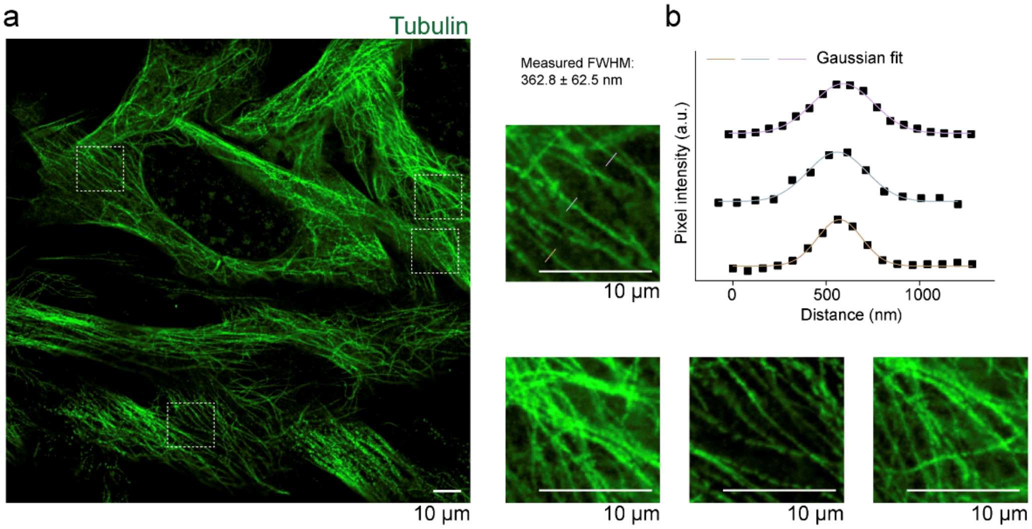


**Figure S1.** Expansion characterization. a. Typical ExM results of C2C12 tubulin expansion. b. Quantification of the line width of single tubulin filaments after expansion.

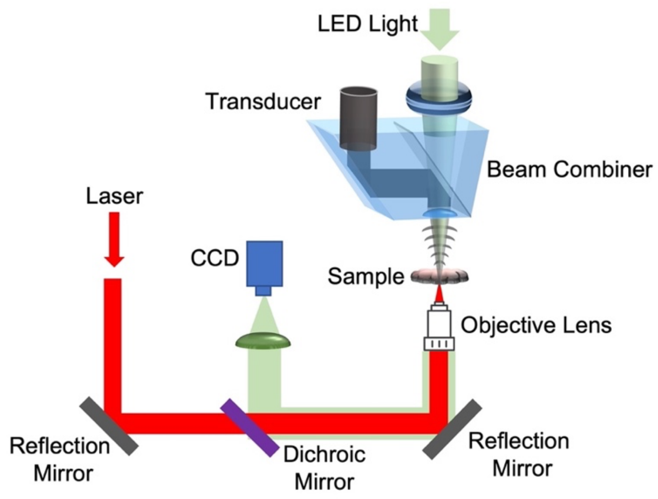


**Figure S2.** Schematic of TOR-PAM.

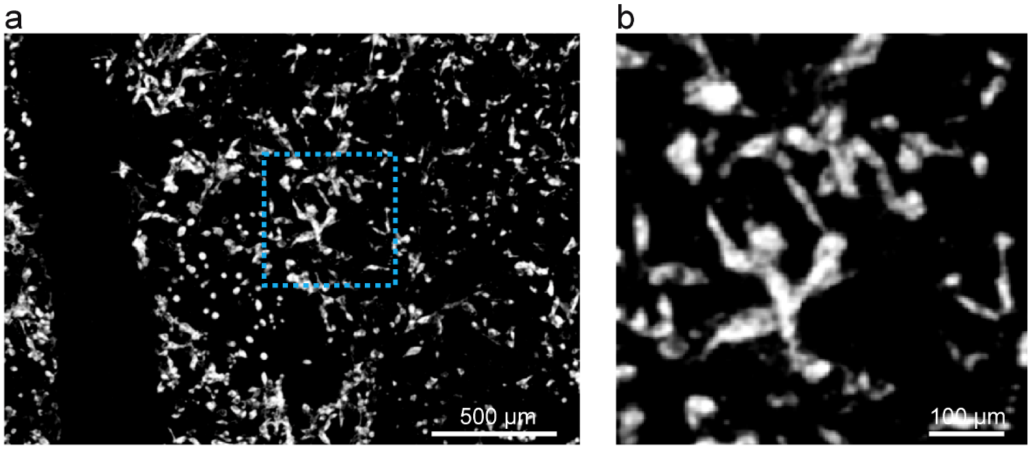


**Figure S3.** a. Photoacoustic images for Vector Blue-stained sample after shrinking back in 10× PBS. b. Magnified image showing the selected area indicated by the blue box in (a).

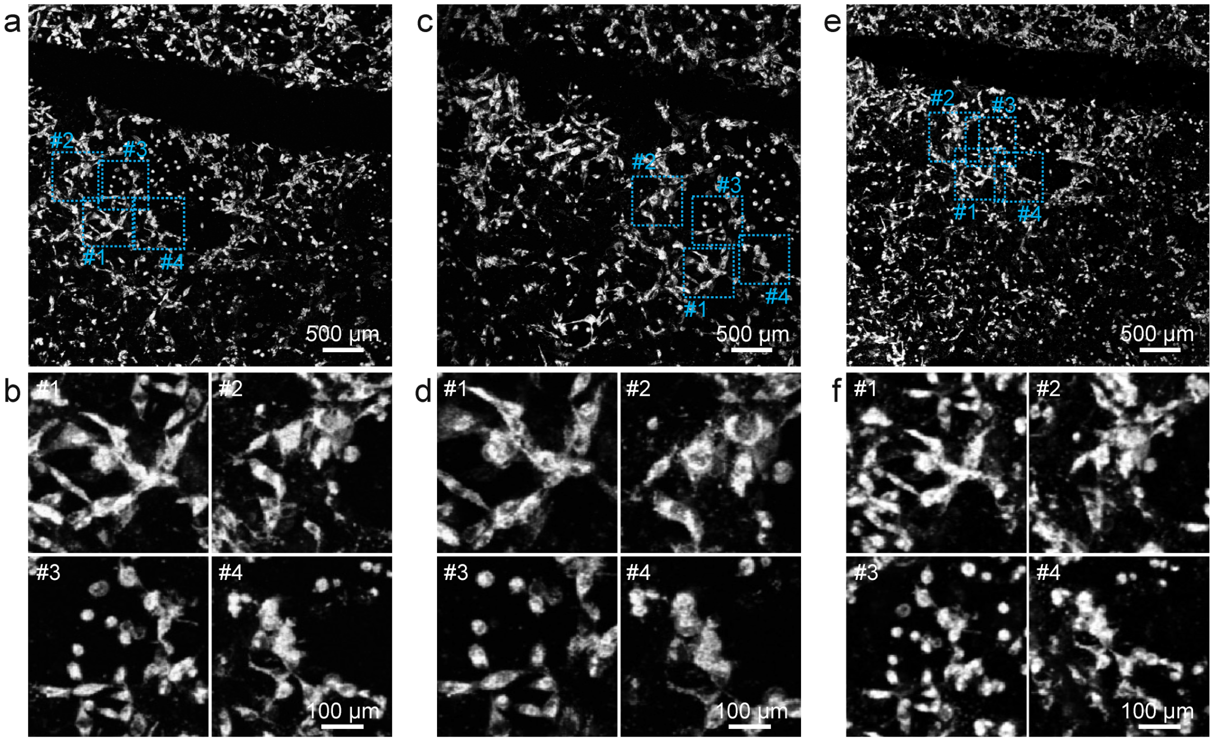


**Figure S4.** PA images of Vector Blue-stained samples. a. Gel before swelling. c. Gel after swelling in water. e. Gel shrinking back in 10× PBS. b, d, and f show magnified areas #1, #2, #3, and #4.

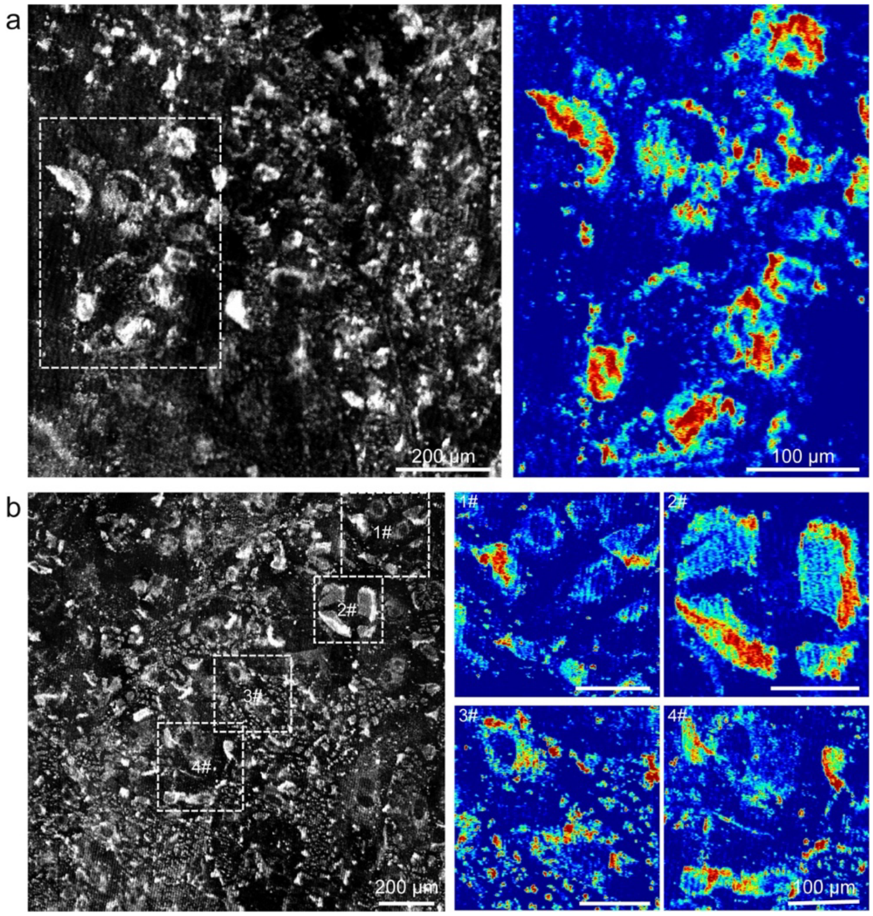


**Figure S5.** Photoacoustic images of the swollen NovaRED-stained sample. a and b show two different areas.

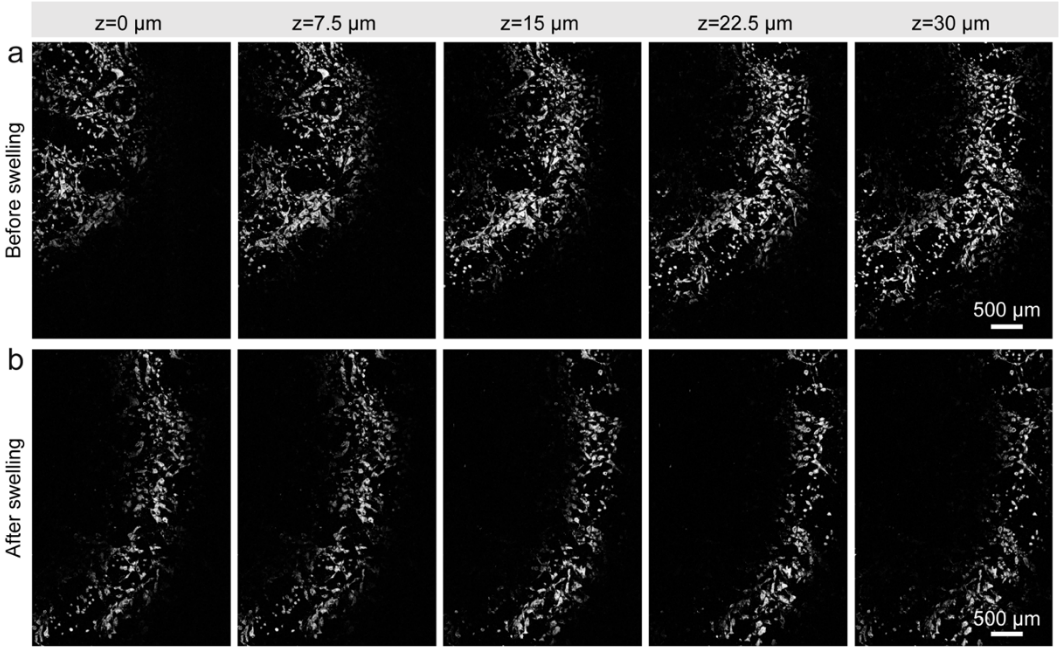


**Figure S6.** Photoacoustic images of the Vector Blue-stained sample showing different depths. a. Gel before swelling. b. Gel after swelling in water.
